# Origin and evolution of the self-organizing cytoskeleton in the network of eukaryotic organelles

**DOI:** 10.1101/005868

**Authors:** Gáspár Jékely

## Abstract

The eukaryotic cytoskeleton evolved from prokaryotic cytomotive filaments. Prokaryotic filament systems show bewildering structural and dynamic complexity, and in many aspects prefigure the self-organizing properties of the eukaryotic cytoskeleton. Here I compare the dynamic properties of the prokaryotic and eukaryotic cytoskeleton, and discuss how these relate to function and the evolution of organellar networks. The evolution of new aspects of filament dynamics in eukaryotes, including severing and branching, and the advent of molecular motors converted the eukaryotic cytoskeleton into a self-organizing ‘active gel’, the dynamics of which can only be described with computational models. Advances in modeling and comparative genomics hold promise of a better understanding of the evolution of the self-organizing cytoskeleton in early eukaryotes, and its role in the evolution of novel eukaryotic functions, such as amoeboid motility, mitosis, and ciliary swimming.

## Introduction

The eukaryotic cytoskeleton organizes space on the cellular scale, and this organization influences almost every process in the cell. Organization depends on the mechano-chemical properties of the cytoskeleton that dynamically maintain cell shape, position organelles and macromolecules by trafficking, and drive locomotion via actin-rich cellular protrusions, ciliary beating or ciliary gliding. The eukaryotic cytoskeleton is best described as an ‘active gel’, a cross-linked network of polymers (gel), where many of the links are active motors that can move the polymers relative to each other (Karsenti et al. 2006).

Since prokaryotes have only cytoskeletal polymers but lack motor proteins, this ‘active gel’ property clearly sets the eukaryotic cytoskeleton apart from prokaryotic filament systems. Prokaryotes contain elaborate systems of several cytomotive filaments (Löwe and Amos 2009) that share many structural and dynamic features with eukaryotic actin filaments and microtubules (Löwe and Amos 1998; van den Ent et al. 2001). Prokaryotic cytoskeletal filaments may trace back to the first cells, and may have originated as higher-order assemblies of enzymes (Noree et al. 2010; Barry and Gitai 2011). These cytomotive filaments are required for the segregation of low copy number plasmids, for cell rigidity and cell wall synthesis, for cell division, and occasionally for the organization of membranous organelles (Thanbichler and Shapiro 2008; Löwe and Amos 2009; Komeili et al. 2006). These functions are performed by dynamic filament-forming systems that harness the energy from nucleotide hydrolysis to generate forces either via bending or polymerization (Löwe and Amos 2009; Pilhofer and Jensen 2013). Although the identification of actin and tubulin homologs in prokaryotes is a major breakthrough, we are far from understanding the origin of the structural and dynamic complexity of the eukaryotic cytoskeleton. Advances in genome sequencing and comparative genomics now allow a detailed reconstruction of the cytoskeletal components present in the last common ancestor of eukaryotes. These studies all point to an ancestrally complex cytoskeleton, with several families of motors (Wickstead et al. 2010; Wickstead and Gull 2007), and filament-associated proteins and other regulators in place (Eme et al. 2009; Fritz-Laylin et al. 2010; Richards and Cavalier-Smith 2005; Jékely 2003; Chalkia et al. 2008; Rivero and Cvrcková 2007; Hammesfahr and Kollmar 2012; Eckert et al. 2011). Genomic reconstructions and comparative cell biology of single-celled eukaryotes (Raikov 1994; Cavalier-Smith 2013) allows us to infer the cellular features of the ancestral eukaryote. These analyses indicate that amoeboid motility (Fritz-Laylin et al. 2010) (although see (Cavalier-Smith 2013)), cilia (Cavalier-Smith 2002; Jékely and Arendt 2006; Mitchell 2004; Satir et al. 2008), centrioles (Carvalho-Santos et al. 2010), phagocytosis (Cavalier-Smith 2002; Jékely 2007; Yutin et al. 2009), a midbody during cell division (Eme et al. 2009), mitosis (Raikov 1994), and meiosis (Ramesh et al. 2005) were all ancestral eukaryotic cellular features. The availability of functional information from organisms other than animals and yeasts (e.g. *Chlamydomonas*, *Tetrahymena, Trypanosoma*) also allow more reliable inferences about the ancestral functions of cytoskeletal components (i.e. not only their ancestral presence or absence) and their regulation (Suryavanshi et al. 2010; Demonchy et al. 2009; Lechtreck et al. 2009).

The ancestral complexity of the cytoskeleton in eukaryotes leaves a huge gap between prokaryotes and the earliest eukaryote we can reconstruct (provided that our rooting of the tree is correct (Cavalier-Smith 2013)). Nevertheless, we can attempt to infer the series of events that happened along the stem lineage, leading to the last common ancestor of eukaryotes. Meaningful answers will require the use of a combination of gene family history reconstructions (Wickstead et al. 2010; Wickstead and Gull 2007), transition analyses (Cavalier-Smith 2002), and computer simulations relevant to cell evolution (Jékely 2008).

## Overview of cytoskeletal functions in prokaryotes and eukaryotes

In the first section I provide an overview of the functions and components of the cytoskeleton in prokaryotes and eukaryotes. To obtain a general overview, I represented cellular structures (e.g. cell wall, kinetochore) and cytoskeletal proteins of prokaryotes and eukaryotes as networks (Figs. 1, 2). In the networks, the nodes represent proteins or cellular structures, and the edges represent the co-occurrence of terms in PubMed entries, used as a proxy for functional connections. The nodes are clustered based on an attractive force (Frickey and Lupas 2004), calculated as the number of entries where the two terms co-occur divided by the number of entries in which the less frequent term occurs.

**Figure 1.**
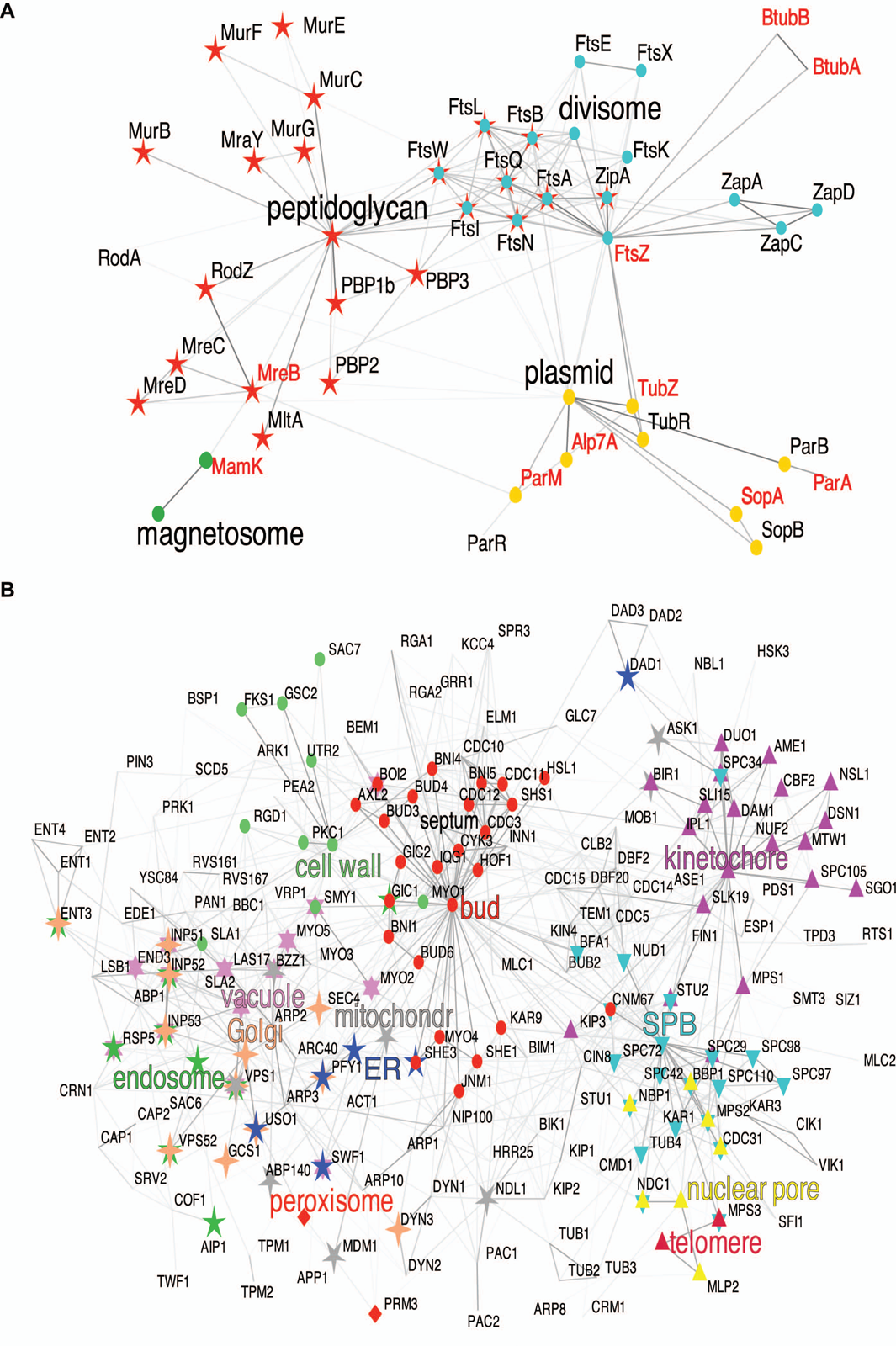
Prokaryotic and yeast cytoskeletal-organellar network. Cytoskeletal-organellar network of (A) prokaryotes and (B) yeast. The nodes correspond to gene names or cytological terms.

**Figure 2.**
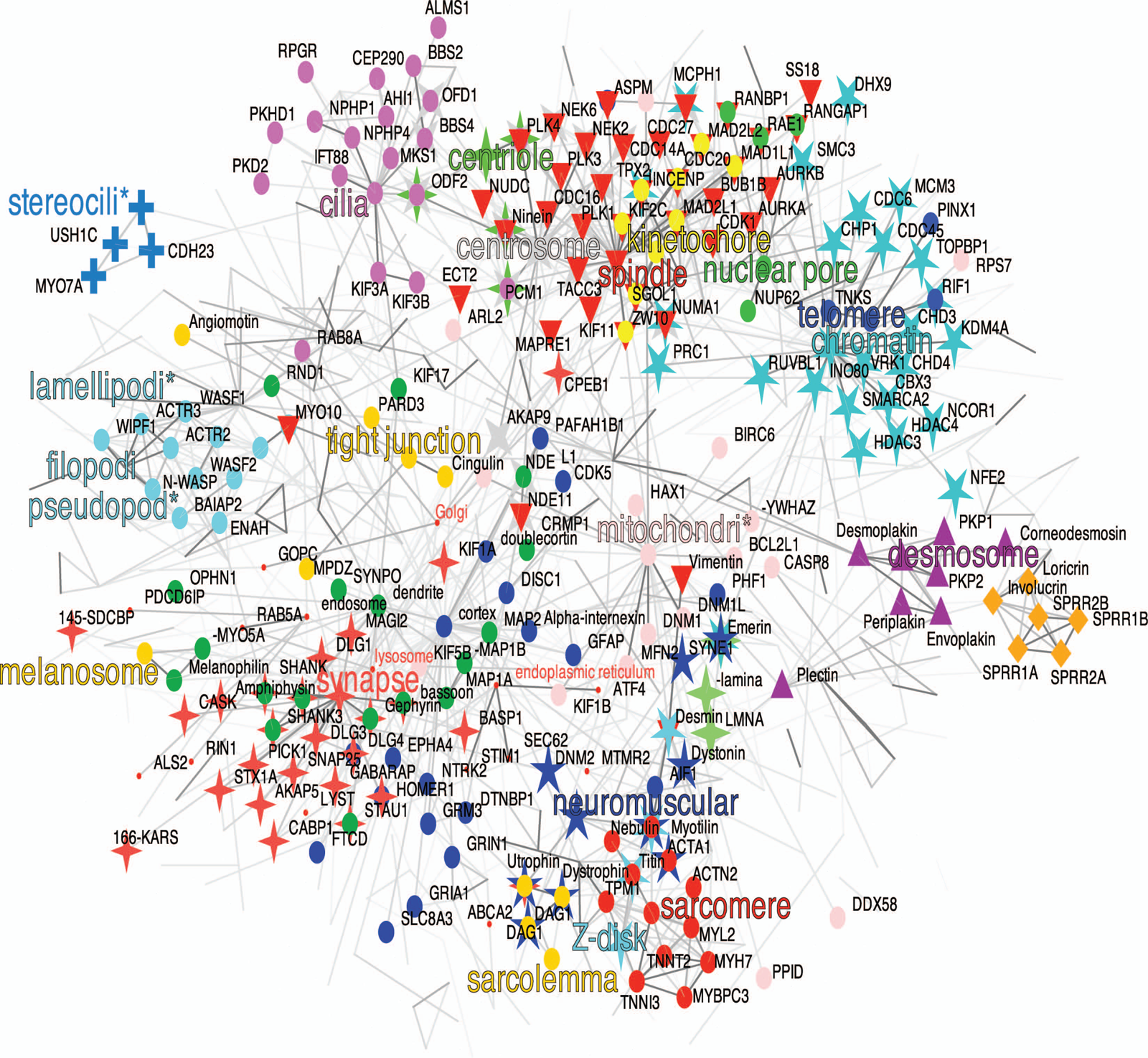
Human cytoskeletal-organellar network. The nodes correspond to gene names or cytological terms.

For prokaryotes, I represented all filament types in one map, even though many of these are specific to certain taxa and do not coexist in the same cell (Fig. 1A). For eukaryotes, I depicted the budding yeast (*Saccharomyces cerevisiae*) cytoskeletal network (Fig. 1B) and a simplified human cytoskeletal network (Fig. 2; eukaryotic cytoskeletal proteins were retrieved from Uniprot using the GO ID GO:0005856).

The prokaryotic network has three major modules, the plasmid partitioning systems, the cell division machinery (divisome) employing the FtsZ contractile ring, and the MreB filament system involved in cell wall synthesis and scaffolding.

Components of the first prokaryotic cytoskeletal module function in the positioning of DNA within the cell, driven by forces generated either by the polymerization or the depolymerization of filaments. These widespread and diverse filament systems are either responsible for the segregation of low copy number plasmids, or for chromosome segregation (Pilhofer and Jensen 2013). DNA partitioning systems generally consist of a centromere-like region on DNA, a DNA-binding adaptor protein, and a filament-forming NTPase, that polymerizes in a nucleotide-dependent manner. Three types of filament systems have been described in prokaryotes. Type I systems employ Walker ATPases (ParA-like), type II systems have actin-like ATPases (ParM-like), and type III systems have tubulin-like GTPases (TubZ-like).

The second widespread prokaryotic filament system functions in cell division. Cell division in all eubacteria and most archaebacteria relies on FtsZ-mediated binary fission. The tubulin-like GTPase, FtsZ (Löwe and Amos 1998), forms filaments that organize into a contractile ring (‘Z-ring’) at the cell centre and trigger fission. The Z-ring is thought to be attached to the membrane at the division site by an ‘A-ring’, formed by the actin-like filament-forming protein, FtsA (Szwedziak et al. 2012). GTP-dependent FtsZ-filament bending may initiate membrane constriction (Osawa et al. 2009). The Z-ring also recruits several downstream components (e.g. FtsI, FtsW) that contribute to the remodeling of the peptidoglycan cell-wall during septation (Lutkenhaus et al. 2012). In archaebacteria, that lack a peptidoglycan wall and FtsA (bar one exception), cell division proceeds using a distinct, poorly understood machinery (Makarova et al. 2010).

The third prokaryotic filament system employs MreB, a homolog of actin that can form filaments in an ATP- or GTP-dependent manner (van den Ent et al. 2001). MreB is found in non-spherical bacteria, and is involved in cell-shape maintenance by localizing cell wall synthesis enzymes. MreB is linked to the peptidoglycan precursor synthesis complex (Mur proteins and MraY) and the peptidoglycan assembly complex (PBPs and lytic enzymes e.g. MltA). Loss of MreB leads to the growth of large, malformed cells that show membrane invaginations (Bendezu and de Boer 2008). *In vitro*, MreB forms filament bundles and sheets (Popp et al. 2010c), whereas *in vivo* MreB filaments form patches under the inner membrane that move together with the cell wall synthesis machinery, probably driven by peptidoglycan synthesis (Domínguez-Escobar et al. 2011; Garner et al. 2011). MreB filament patches also contribute to the mechanical rigidity of the cell, independent of their function in cell wall synthesis (Wang et al. 2010).

The eukaryotic cytoskeletal networks (represented by yeast and human) include a cell-division module including the spindle, centromere, and the centrosome (spindle pole body, SPB in yeast). This module functions in chromosome segregation, during which kinetochores must interact with spindle microtubules. Proper attachment is for example facilitated by Stu2 (ortholog of vertebrate XMAP215), a protein that is transferred to shrinking microtubule plus ends when they reach a kinetochore, and stabilizes them (Gandhi et al. 2011). Other examples from this module are Aurora kinase and INCENP (yeast Ipl1 and Sli15), proteins that ensure that sister kinetochores attach to microtubules from opposite spindle poles during mitosis (Tanaka et al. 2002).

Another important subnetwork in the yeast cytoskeleton is involved in bud-site selection and the formation of a contractile actomyosin ring. An example in this network is yeast Myo1, a two-headed myosin-II that localizes to the division site and promotes the assembly of a contractile actomyosin ring and septum formation (Fang et al. 2010). The membrane trafficking subnetwork includes regulators of vesicle trafficking and cargo sorting, including the yeast dynamin-like GTPase, Vps1. Vps1 is involved in vacuolar, Golgi and endocytic trafficking (Vater et al. 1992).

The human cytoskeletal network includes several other modules absent from yeast. These include a module centered around the cilium, and one module for the formation of lamellipodia, filopodia, and phagocytosis. The former includes ciliary transport (intraflagellar transport, BBSome), structural, and signaling (PKD2) proteins, the latter includes proteins that reorganize cortical actin filaments, including the Arp2/3 complex (ACTR2/3 (Mullins et al. 1998)) and the Cdc42 effector N-WASP, an activator of the Arp2/3 complex (Takenawa and Miki 2001). The human network also contains several animal-specific modules, including modules related to stereocilia of inner-ear hair-cells, muscle, neurons (dendrite, synapse), skin, and structures mediating cell-cell adhesion (desmosome).

Despite the vastly different organization and complexity of the eukaryotic and prokaryotic cytoskeletal networks, we know that there is evolutionary continuity between them. The eukaryotic cytoskeletal networks are centered around actin-like and microtubule-like cytomotive filaments, that evolved from homologous filament systems in prokaryotes (Löwe and Amos 1998; van den Ent et al. 2001).

## Prokaryotic origin of the major components of the eukaryotic cytoskeleton

In this section I give an overview of the diversity of actin- and tubulin-like filament-forming proteins, and discuss a few other key cytoskeletal components, for which distant prokaryotic homologs could be identified.

Besides actin- and tubulin-like filaments, prokaryotes also contain filament-forming Walker ATPases (ParA and SopA), with no polymer-forming homologs in eukaryotes. The evolution of this family will not be discussed.

### Origin of eukaryotic actin filaments

Actin is a member of the sugar kinase/HSP70/actin superfamily (Bork et al. 1992). This family also includes different prokaryotic filament-forming proteins, including MreB, FtsA, the plasmid-partitioning protein ParM and its relatives, and an actin family specific to archaebacteria (crenactins).

To represent the diversity of actin-like proteins and their phyletic distribution in a global map, I clustered a large dataset of actin-like sequences based on pairwise BLASTP *P* values using force-field based clustering (Frickey and Lupas 2004) (Fig. 3 A-C). Clustering can be very efficient if large numbers of sequences need to be analyzed. Given that, at least in prokaryotes, there is a tight link between orthologs and bidirectional best BLAST hits (Wolf and Koonin 2012), BLAST-based clustering can efficiently recover orthology groups in large datasets. Even though clustering methods still lack sophisticated analysis tools that are common in alignment-based molecular phylogeny methods (e.g. rate heterogeneity among sites), the results from similarity-based clustering can agree well with molecular phylogeny (Jékely 2013; Mirabeau and Joly 2013). Cluster maps can also provide a general overview of taxonomic distribution and of sequence similarity, parameters that are not easily inferred from phylogenetic trees. Clustering is best though of as a representation of sequence data as a similarity network, allowing evolutionary biologists to draw inferences about sequence evolution than are complementary to answers based on phylogeny (for a thoughtful introduction to the use of similarity networks see (Halary et al. 2013)).

**Figure 3.**
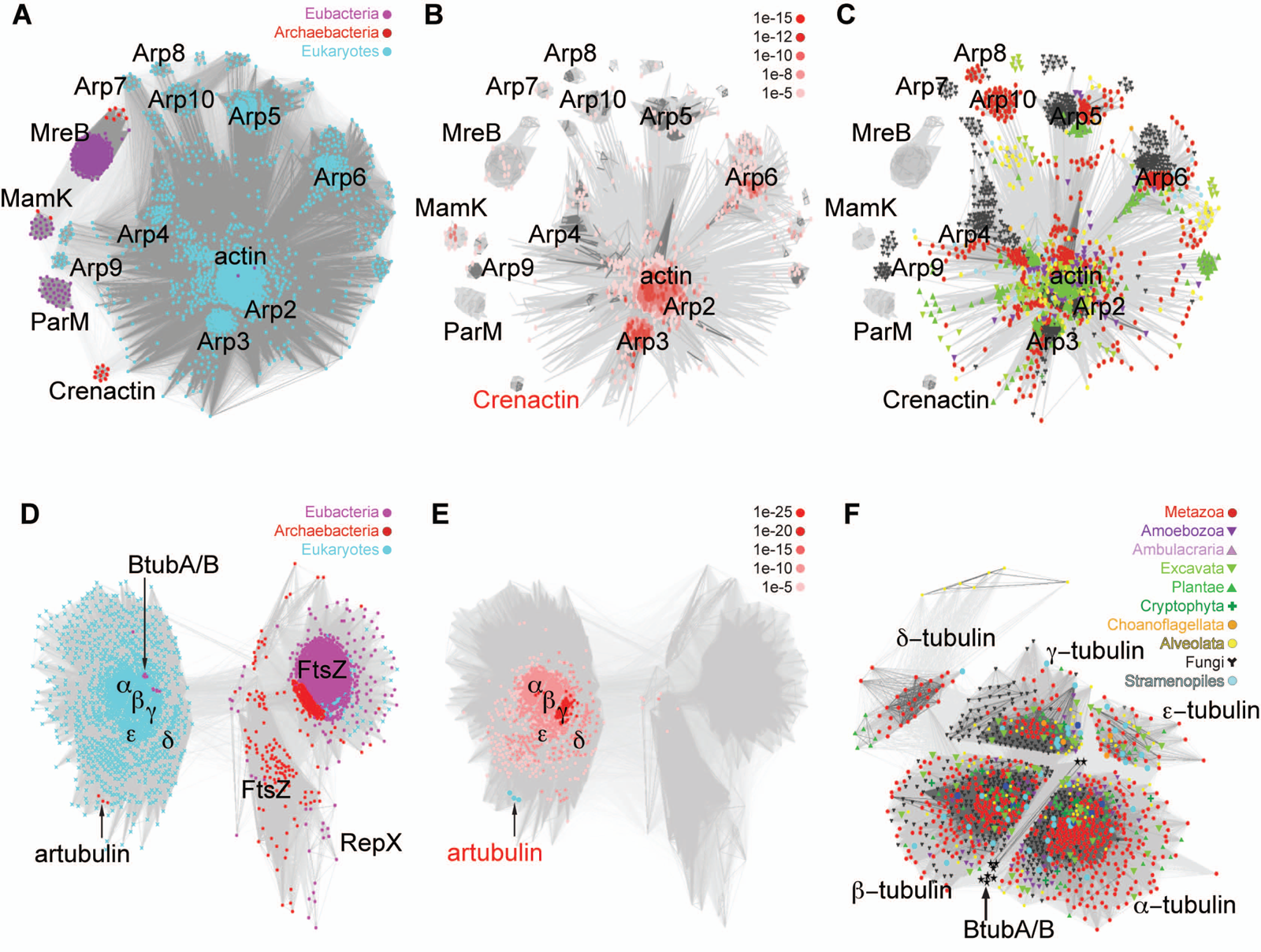
Cluster analysis of actin-like and tubulin-like proteins. Sequence-similarity-based clustering was performed on (A-C) prokaryotic and eukaryotic actin-like proteins, and (D-F) prokaryotic and eukaryotic tubulin-like proteins. In both cases an exhaustive, 90% non-redundant set of Uniprot is shown. The clusters were colored to reflect domain-wide (A, D) or eukaryote-wide (C, F) phyletic distribution. The BLASTP connections of (B) crenactins and (E) artubulins were shown, with hits of different *P* value cutoffs shown in different hues of red.

The actin similarity network revealed all actin-like protein families and their phyletic distribution. Filamentous actin was at the centre of the cluster of eukaryotic actins, and the diverse Arp families radiated from this centre. The ‘centroid’ position of actin (Fritz-Laylin et al. 2010) suggests that it represents the most ancestral eukaryotic sequence, and therefore maximizes all the blast hits to other eukaryotic actins. The ancestral nature of actin is also in agreement with its role in filament formation, whereas the more derived Arps are either regulators of filament branching and nucleation (the Arp2/3 complex (Mullins et al. 1998)), or have unrelated functions.

The similarity map also reveals the prokaryotic MreB, MamK, ParM, and crenactin families (the more derived FtsA was excluded) as distinct clusters. Among the prokaryotic actins, crenactins show the most similarity to eukaryotic actins, and have been proposed to be the direct ancestors of eukaryotic actins (Bernander et al. 2011; Yutin et al. 2009). Crenactin was shown to form helical structures in *Pyrobaculum* cells, and is only found in rod-shaped archaebacteria (Ettema et al. 2011), indicating that it may regulate cell shape. Crenactins share two unique inserts with eukaryotic actins, and other inserts that are uniquely shared with the actin-like protein Arp3 (Yutin et al. 2009). This is a puzzling observation, and either suggest that Arp3 (arguably a derived regulatory actin) represents the ancestral state, or that crenactins originated via horizontal gene transfer (HGT) from eukaryotes to archaebacteria. The phylogenetic trees showing a sister relationship of crenactins to all eukaryotic actins (Rolf Bernander 2011; Yutin et al. 2009) should be interpreted with caution, given that these trees have long internal branches, use very distant outgroups, and have few aligned positions. If crenactins were derived Arp3 proteins, they would also be expected to branch artificially at a deeper node, not as a sister to Arp3, due to long-branch attraction. Future structural studies of crenactins may be able to clarify the history of crenactins, relative to eukaryotic actins.

### Origin of microtubules

Microtubules are dynamic polymer tubes formed by 13 laterally interacting protofilaments of α/β-tubulin heterodimers. Like actin filaments, microtubules are universal in eukaryotes. Besides the canonical α/β-tubulins, several other tubulin forms have ancestrally been present in eukaryotes, including delta, gamma and epsilon tubulins. The prokaryotic homologs of tubulins include FtsZ, TubA, BtubA/BtubB from Verrucomicrobia, and artubulins, so far only found in the archaebacterium *Nitrosoarchaeum* (Yutin and Koonin 2012).

The cluster map of tubulins provides an overview of the phyletic distribution of all families (Fig. 3 D-F). Alpha, beta, gamma, delta, and epsilon tubulins are all ancestrally present in eukaryotes, given their broad distribution and their presence in excavates, a protist group that potentially represents a divergence close to the root of the eukaryotic tree (Cavalier-Smith 2013). Epsilon and delta tubulin are only present in lineages with a cilium.

There are two independent, phyletically restricted groups of prokaryotic tubulins with higher sequence similarity to eukaryotic tubulins than FtsZ, BtubA/BtubB from *Prosthecobacter* and the archaebacterial artubulins.

BtubA and BtubB were identified in *Prosthecobacter* (Jenkins et al. 2002), belonging to the Verrucomicrobia. These proteins show high sequence (∼35% identity) and structural similarity to eukaryotic α/β-tubulins, and form tubulin-like protofilaments, made up of BtubA/BtubB heterodimers (Schlieper et al. 2005). Despite the close similarity to α/β-tubulins, there is no one-to-one correspondence between the α/β and BtubA/BtubB heterodimers. Instead, both BtubA and BtubB exhibit structural features that are specific to either α or β tubulin (Schlieper et al. 2005). This, together with the equal distance from α/β-tubulin in sequence space (Fig. 3F), suggests that BtubA/BtubB represent a state in tubulin evolution preceding the duplication of α/β-tubulins in stem eukaryotes. Since α/β-tubulins are the structural components of microtubules, their origin by gene duplication was probably the first event in the history of eukaryotic tubulin duplications. The close similarity of BtubA/BtubB to eukaryotic tubulins suggests that they originated by HGT from eukaryotes to *Prosthecobacter* (Schlieper et al. 2005).

Nevertheless, the ancestral character of BtubA/BtubB, uniting features of α/β-tubulin suggests that BtubA/BtubB originated by an ancient HGT event, and these tubulins may provide insights into the early evolution of microtubules. Interestingly, and in contrast to all other prokaryotic tubulins, BtubA/BtubB can form tubules formed by 5 protofilaments (instead of 13 as in eukaryotes) (Pilhofer et al. 2011). These simpler, smaller tubules may represent an intermediate stage in the evolution of the eukaryotic tubulin skeleton. The ability to form microtubules may also explain the higher sequence conservation of BtubA/BtubB, despite their potential early origin.

Another class of prokaryotic tubulins, artubulin, has recently been identified in *Nitrosoarchaeum* and has been proposed to be the ancestors of eukaryotic tubulins (Yutin and Koonin 2012). Artubulins show higher sequence similarity to eukaryotic tubulins, than to FtsZ. In a phylogenetic tree artubulins branched as a sister to all eukaryotic tubulins. In the cluster map, artubulins appear at the periphery of the eukaryotic tubulins (Fig. 3D), and show very low sequence similarity to FtsZ. Coloring the nodes connected to artubulins according to their similarity (BLAST P value) to artubulins indicates that gamma-tubulins are closest in sequence space. Gamma-tubulin regulates microtubule nucleation, and it is more likely that it represents a derived tubulin class, not one ancestral to α/β-tubulins. These considerations, together with the very limited taxonomic distribution of artubulins cast further doubt on their ancestral status. The clustering results are consistent with artubulins representing a derived gamma tubulin, acquired by HGT from eukaryotes to *Nitrosoarchaeum*. The original molecular phylogeny may have grouped artubulins deep due to a long-branch artifact, caused both by the derived nature of artubulins and the very distant outgroup. Structural analysis and polymerization assays of artubulins will help to better evaluate these alternative scenarios.

Overall, the origin of eukaryotic tubulin from either BtubA/BtubB or artubulins is not convincingly demonstrated, and both may have been acquired by HGT from eukaryotes. If this is the case, then the most likely ancestor of eukaryotic tubulins remains to be FtsZ.

### Origin of molecular motors

Molecular motors are mechano-chemical enzymes that use ATP hydrolysis to drive a mechanical cycle (Vale and Milligan 2000). Motors step either along microtubules (kinesins and dyneins) or the actin cytoskeleton (myosins) and are linked to and move cargo (molecules or organelles) around the cell. Several families of all three motor types are ancestrally present in eukaryotes (Wickstead and Gull 2007; 2011; Wickstead et al. 2010; Richards and Cavalier-Smith 2005).

The origin of motors is unknown, since no direct prokaryotic ancestor has been identified. However, kinesins and myosins have common ancestry and share a catalytic core and a ‘relay helix’ that transmits the conformational change in the catalytic core to the polymer binding sites and the mechanical elements (Jon Kull et al. 1996). These motors are distantly related and evolved from GTPase switches (Leipe et al. 2002), molecules that likewise undergo conformational changes upon nucleotide binding and hydrolysis (Vale and Milligan 2000).

### Other prokaryotic homologs of cytoskeletal proteins

The prokaryotic ancestry of cytoskeletal components other than actin and tubulin can also be ascertained by sensitive sequence and structural comparisons.

Profilin is a protein that breaks actin filaments (Schutt et al. 1993). Profile-profile searches with profilin using HHpred recovered the bacterial gliding protein MglB (Probab = 95.44 E-value = 0.73) and other proteins with the related Roadblock/LC7 domain. Profilin and MglB also show structural similarity, as shown by PDBeFold searches (profilin 3d9y:B and MglB 3t1q:B with an RMSD of 2.65). The sequence- and structure-based similarities establish profilin as a homolog of the Roadblock family. In eukaryotes, members of this family are associated with ciliary and cytoplasmic dynein, and in prokaryotes MglB is a GTPase activating protein (GAP) of the gliding protein, the Ras-like GTPase MglA (Leonardy et al. 2010).

The microtubule severing factors katanin and spastin, members of the AAA+ ATPase family, also have prokaryotic origin. AAA+ ATPases have several ancient families with broad phyletic distribution, and the katanin family is a member of the classical AAA clade (Iyer et al. 2004). This clade includes bacterial FtsH (an AAA+ ATPase with a C-terminal metalloprotease domain), a protein that is localized to the septum in dividing *Bacillus subtilis* cells (Wehrl et al. 2000) where it may degrade FtsZ (Anilkumar et al. 2001). Whether the katanin family of microtubule severing factors evolved by the modification of FtsH, is not resolved.

A third cytoskeletal regulator with prokaryotic ancestry is the enzyme alpha-tubulin N-acetyltransferase (mec-17) that stabilizes microtubules in cilia and neurites by alpha-tubulin acetylation. The phyletic distribution of this enzyme tightly parallels that of cilia. Alpha-tubulin N-acetyltransferase is a member of the Gcn5-related N-acetyl-transferase (GNAT) superfamily (Steczkiewicz et al. 2006; Taschner et al. 2012), widespread in prokaryotes (Neuwald and Landsman 1997).

## Dynamic properties of the prokaryotic and eukaryotic cytoskeleton

In the following section I compare the dynamic properties of the prokaryotic and eukaryotic cytoskeleton. Prokaryotic filament-forming systems show remarkable properties that in many respects prefigure the dynamic, self-organized properties of the eukaryotic cytoskeleton (Fig. 4). The dynamic features of prokaryotic filaments include regulated filament nucleation (Lim et al. 2005), polymerization and depolymerization, dynamic instability (Garner 2004), treadmilling (Larsen et al. 2007), directional polarization with plus and minus ends (Larsen et al. 2007), the formation of higher-order structures (Szwedziak et al. 2012), and force-generation by filament growth, shrinkage or bending. These features enable prokaryotic filaments to perform various functions, such as the positioning of membraneous organelles, chromosome and plasmid segregation, cell-shape changes, cell division, and contribution to the mechanical integrity of the cell (Wang et al. 2010).

**Figure 4.**
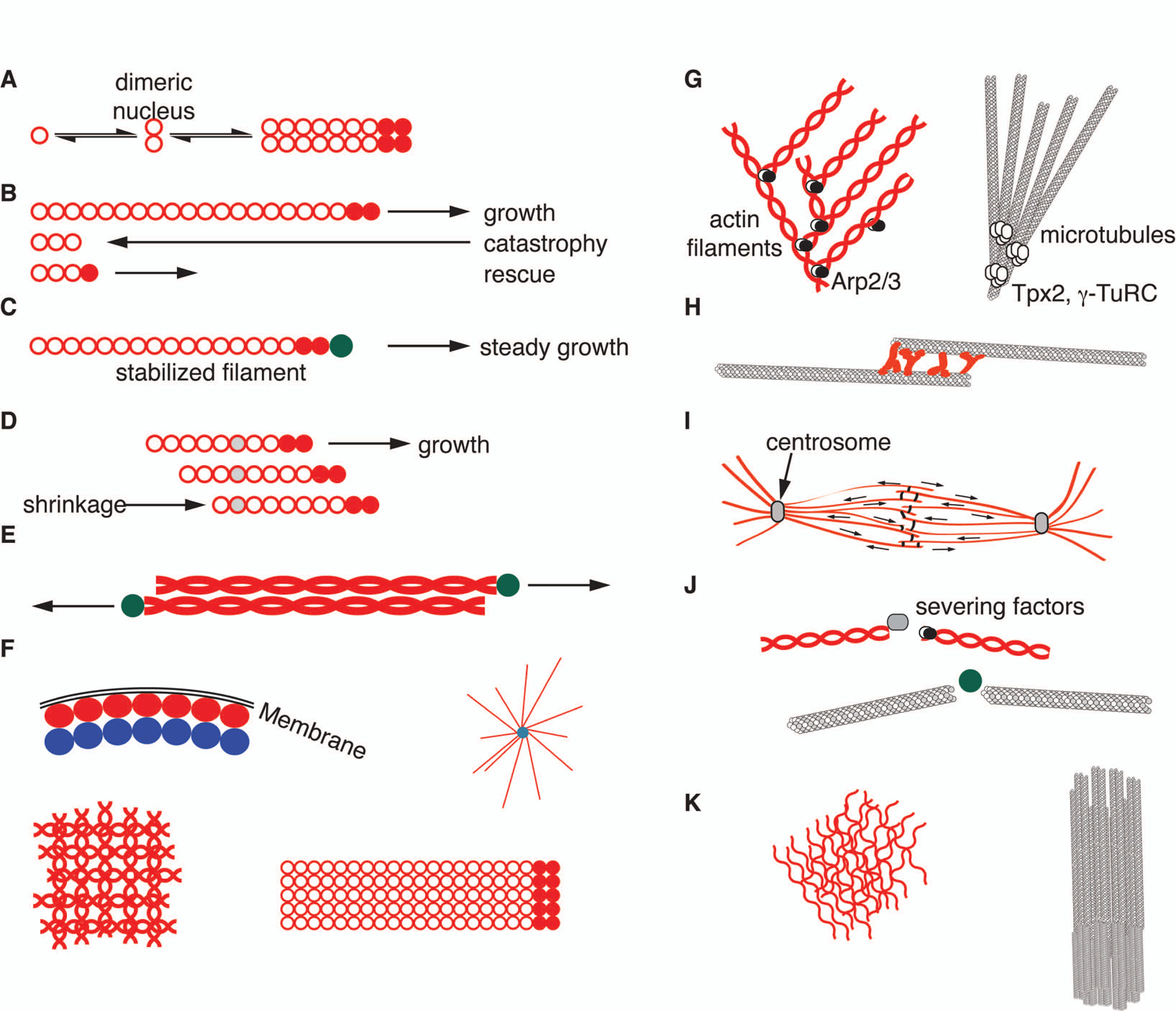
Dynamic properties of the cytoskeleton. Dynamic properties and self-organized patterns of the prokaryotic (A-F) and eukaryotic (G-K) cytoskeleton. (A) Filament nucleation by a dimeric nucleus, (B) dynamic instability, (C) filament capping, (D) treadmilling, (E) bipolar growth of antiparallel filaments, (F) higher-order structures, such as filament pairs, asters, meshes, sheets. Eukaryotes in addition display (G) filament branching, (H) dynamic overlap of antiparallel filaments, (I) spindle and asters, (J) filament severing, (K) actin networks, axoneme and basal bodies.

The eukaryotic cytoskeleton shares all of the above features with the cytomotive filaments of prokaryotes (Fig. 4) but evolved additional features (Table 1). First I discuss the dynamic properties shared between prokaryotes and eukaryotes. I then give an overview of the unique properties of the eukaryotic cytoskeleton that represent evolutionary innovations during the origin of eukaryotes.

**Table 1.**
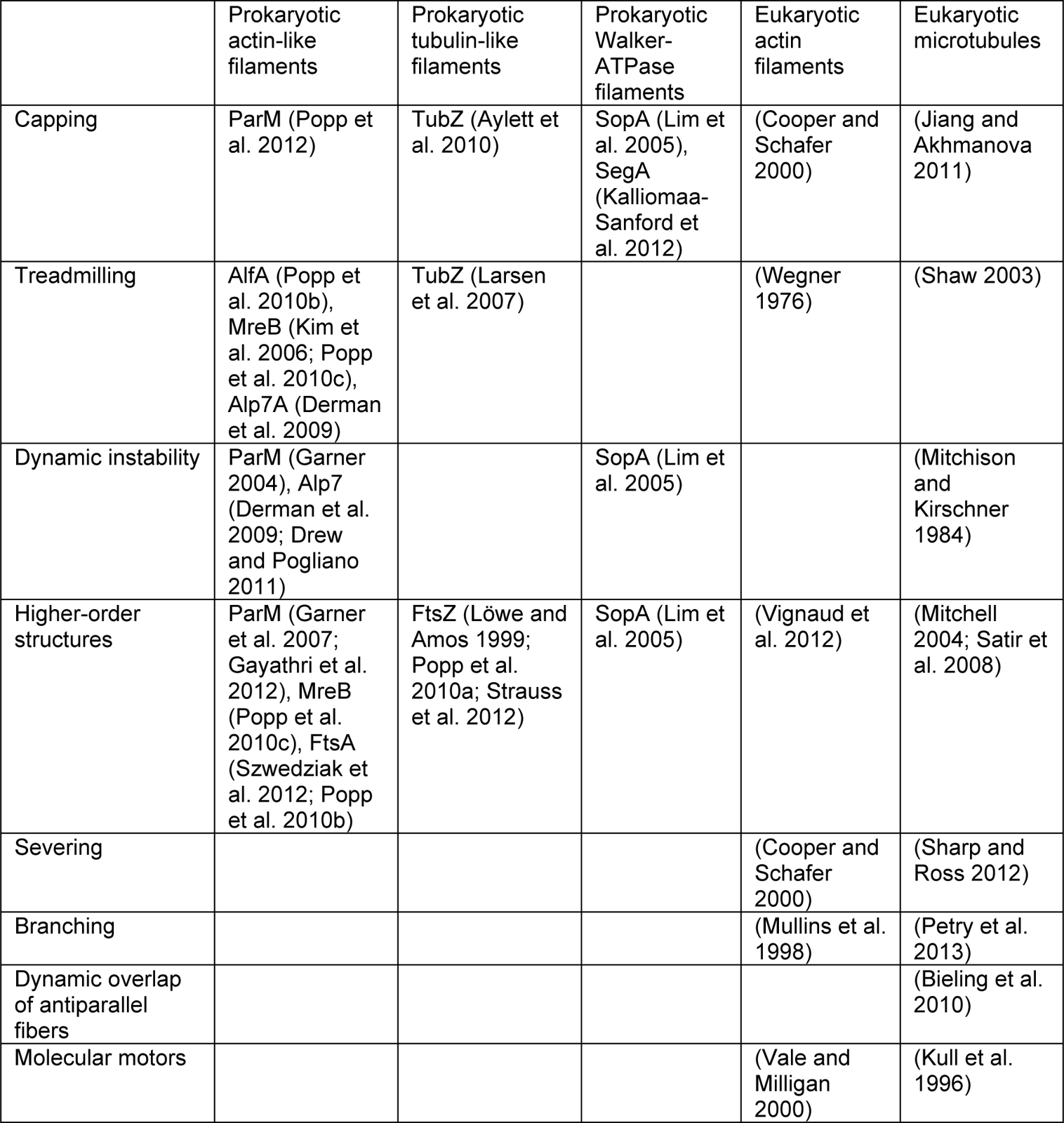
Overview of the dynamic properties of prokaryotic and eukaryotic filament systems.

|  | Prokaryotic actin-like filaments | Prokaryotic tubulin-like filaments | Prokaryotic Walker-ATPase filaments | Eukaryotic actin filaments | Eukaryotic microtubules |
| --- | --- | --- | --- | --- | --- |
| Capping | ParM (Popp et al. 2012) | TubZ (Aylett et al. 2010) | SopA (Lim et al. 2005), SegA (Kalliomaa-Sanford et al. 2012) | (Cooper and Schafer 2000) | (Jiang and Akhmanova 2011) |
| Treadmilling | AlfA (Popp et al. 2010b), MreB (Kim et al. 2006; Popp et al. 2010c), Alp7A (Derman et al. 2009) | TubZ (Larsen et al. 2007) |  | (Wegner 1976) | (Shaw 2003) |
| Dynamic instability | ParM (Garner 2004), Alp7 (Derman et al. 2009; Drew and Pogliano 2011) |  | SopA (Lim et al. 2005) |  | (Mitchison and Kirschner 1984) |
| Higher-order structures | ParM (Garner et al. 2007; Gayathri et al. 2012), MreB (Popp et al. 2010c), FtsA (Szwedziak et al. 2012; Popp et al. 2010b) | FtsZ (Löwe and Amos 1999; Popp et al. 2010a; Strauss et al. 2012) | SopA (Lim et al. 2005) | (Vignaud et al. 2012) | (Mitchell 2004; Satir et al. 2008) |
| Severing |  |  |  | (Cooper and Schafer 2000) | (Sharp and Ross 2012) |
| Branching |  |  |  | (Mullins et al. 1998) | (Petry et al. 2013) |
| Dynamic overlap of antiparallel fibers |  |  |  |  | (Bieling et al. 2010) |
| Molecular motors |  |  |  | (Vale and Milligan 2000) | (Kull et al. 1996) |

### Filament nucleation, polymerization, depolymerization, and capping

The regulation of the polymerization and depolymerization of polymers is essential for the proper functioning of filament systems. Filament growth can be influenced by various factors, including monomer concentration, nucleotides, and accessory factors, such as nucleating or polymerizing proteins. In eukaryotes, the spontaneous nucleation of microtubules and actin filaments is slow. Filament nucleation therefore represents an important regulatory component, allowing the positioning of growing filaments in the cell (Goley and Welch 2006; Kollman et al. 2011). In contrast, prokaryotic filaments commonly assemble rapidly and spontaneously, although nucleation may in some cases facilitate assembly. For example, FtsZ filament assembly proceeds via an FtsZ dimer that can serve as a nucleus for polymerization (Chen et al. 2005).

Cytoskeletal filament dynamics is also regulated by filament capping. Capping includes the binding of factors to the end of a filament, thereby preventing disassembly. Eukaryotic actin fibres and microtubules are both regulated by capping (Cooper and Schafer 2000) (Jiang and Akhmanova 2011). In prokaryotic DNA partitioning systems filament assembly is commonly facilitated by the centromere-adaptor protein complex that stabilizes the growing end of the filament. This has been observed for all three types of prokaryotic filaments (Table 1)(Lim et al. 2005; Kalliomaa-Sanford et al. 2012; Popp et al. 2012; Aylett et al. 2010). Capping by the DNA-adaptor complex ensures the steady polymerization of the filaments by the incorporation of new subunits, thereby moving the plasmid or the chromosome (Salje and Löwe 2008; Kalliomaa-Sanford et al. 2012).

### Treadmilling

Treadmilling is an important feature of eukaryotic actin (Wegner 1976) and microtubules (Shaw 2003), and is characterized by filament polymerization at one end and depolymerization at the other end. This results in the apparent motion of the filament, even though the individual subunits stay in place. Treadmilling has also been observed in the actin-like and tubulin-like DNA segregation proteins (Table 1)(Popp et al. 2010b; Kim et al. 2006; Popp et al. 2010c; Derman et al. 2009; Larsen et al. 2007). Treadmilling of the tubulin-like protein TubZ was shown to be important for plasmid stability. TubZ with a mutation in a catalytic residue forms stable filaments that are unable to undergo treadmilling. The introduction of this mutant into the cell leads to the loss of the associated plasmid, highlighting the importance of filament dynamics for proper plasmid segregation (Larsen et al. 2007).

### Dynamic instability

Cytoskeletal filaments often show dynamic instability, characterized by the alternation of steady polymerization and catastrophic shrinkage. This behavior is also characteristic of eukaryotic microtubules (Mitchison and Kirschner 1984). Microtubues are polar, growing at their plus ends by the addition of tubulin heterodimers. Tubulins use GTP for filament assembly, and GTP hydrolysis within the microtubule generates tension that is required for dynamic instability (Karsenti et al. 2006). Dynamic instability represents an efficient strategy to search in space (Holy and Leibler 1994). Dynamic instability is important for proper DNA capture and positioning in both prokaryotes and eukaryotes. The actin-like proteins ParM (Garner 2004), and Alp7 (Derman et al. 2009) were observed to undergo dynamic instability *in vivo*. The nucleotide-bound monomers form a cap that stabilize the filament (Garner 2004), but upon nucleotide hydrolysis, the filament rapidly disassembles.

ParM filaments are polar, but when two filaments associate in an antiparallel fashion, they polarize bidirectionally (Gayathri et al. 2012). The filaments are dynamic and search the cell. Binding of the ParR/parC adaptor/centromere complex to the ends of ParM filaments inhibits dynamic instability, and promotes filament growth. This ‘search and capture’ mechanism allows efficient plasmid segregation by pushing plasmids apart in a bipolar spindle.

The Walker ATPase SopA also forms dynamic filaments, the dynamics of which is important for plasmid segregation, since mutants that form static polymers inhibit segregation (Lim et al. 2005). Filaments formed by the actin-like Alp7A also undergo dynamic instability, and computational modeling and experiments of an artificial system consisting of Alp7A and a plasmid revealed how such dynamic instability can drive the positioning of plasmids either to the cell centre or the cell poles (Drew and Pogliano 2011). This bimodal system is tunable, and cell-centre or cell-pole positioning depends on the parameters of dynamic instability. This simple system illustrates how a dynamic cytoskeletal system of a few components can create spatial inhomogeneity of macromolecules in the cell.

### Force-generation

The eukaryotic cytoskeleton can generate force by at least three distinct mechanisms, filament growth, filament shrinkage (Kueh and Mitchison 2009; McIntosh et al. 2010), or molecular motors walking on filaments (Vale 2003). In prokaryotes, no motor has been found, and force is generated by filament growth, filament shrinkage, or filament bending (FtsZ). Nucleotide-driven filament growth relies on the continuous addition of subunits to the filament end, which can push the attached structures. This is the general mechanism of force generation for all three types of plasmid partitioning systems. Filament shrinkage has also been suggested as a mechanism of force generation during chromosome segregation in *C. crescentus*. The shrinkage of the ParA filament, destabilized by centromere-bound ParB, is thought to move the centromere to the cell poles through a ‘burnt-bridge Brownian ratchet’ mechanism (Ptacin et al. 2010). Filament bending was proposed to exert force during FtsZ-mediated cell division (Osawa et al. 2009). Large filaments formed by the over-expression of the actin-like protein, FtsA, can also bend *E. coli* cells (Szwedziak et al. 2012).

### Cooperation of distinct filament types

In eukaryotes, the actin and tubulin systems often work together, for example at the midbody during cell division, or during endocytosis when cargo vesicles switch from actin-to microtubule-based transport (Soldati and Schliwa 2006). Cooperation of distinct filament types also occurs in prokaryotes. In *C. crescentus* the CtpS filaments and crescentin filaments co-occur at the inner cell curvature and regulate each other. Crescentin recruits CtpS, and CtpS negatively regulates crescentin assembly (Ingerson-Mahar et al. 2010). A two-filament system is also important during FtsZ-mediated cell division. The tubulin-like GTPase forming the constriction ring, FtsZ, is recruited to the membrane by the actin-like protein FtsA. FtsA also forms filaments, and this ability was shown to be important for proper cell division (Szwedziak et al. 2012). Polymerized FtsZ may be attached to the membrane by patches of polymerized, membrane-bound FtsA, localized to the cell division ring. It has recently been found that FtsZ also directly interacts with MreB, and this interaction is required for Z ring contraction and septum synthesis (Fenton and Gerdes 2013).

### Higher-order filament structures

Eukaryotic filament systems generate several higher-order structures, including the ciliary axoneme, the mitotic spindle, microtubule asters, or contractile actin meshworks (Vignaud et al. 2012). Several prokaryotic filaments also form higher-order structures (Table 1). FtsZ can form toroids and multi-stranded helices, consisting of several filaments bundled together (Popp et al. 2010a). Pairs of parallel FtsZ filaments associated in an antiparallel fashion can form sheets (Löwe and Amos 1999). *In vivo*, FtsZ forms discontinuous patches in a bead-like arrangement at the cell division ring, consisting of several filaments (Strauss et al. 2012). SopA is able to form aster-like structures *in vitro*, radiating from its binding partner, SopB, bound to a plasmid containing SopB-recognition-sites (Lim et al. 2005). The actin-like protein ParM is also able to form asters *in vitro* (Garner et al. 2007), and antiparallel filaments in the cell (Gayathri et al. 2012). MreB forms multilayered sheets with diagonally interwoven filaments, or long cables with parallel protofilaments (Popp et al. 2010c). Filaments of the actin-homolog FtsA can also form large bundles when overexpressed in *E. coli*, that can bend the cell and tubulate the membrane (Szwedziak et al. 2012). The bacterial actin AlfA can form 3D-bundles, rafts and nets (Popp et al. 2010b). This list is impressive and illustrates well the versatility of the prokaryotic cytoskeleton. However, the complexity of the higher-order structures formed by the eukaryotic cytoskeleton far surpasses the complexity of these structures. The eukaryotic cytoskeleton organizes cellular space both using dynamic scaffolds (e.g. mitotic spindle, microtubule aster, lamellipodia) and static scaffolds built of stabilized filaments (e.g. axoneme, microtubular ciliary root, microtubule-supported cell-cortex in several protists, stabilized microtubule bundles in metazoan neurites, microvilli, sarcomeres). The formation of these structures would not be possible without the unique dynamic properties of the eukaryotic cytoskeleton.

### Unique dynamic properties of the eukaryotic cytoskeleton

The eukaryotic cytoskeleton has several novel properties, not yet described in prokaryotic filament systems. These include filament severing (actin fibers (Cooper and Schafer 2000) and microtubules (Sharp and Ross 2012)), branching (actin fibers (Mullins et al. 1998) and microtubules (Petry et al. 2013)), and dynamic overlap of the antiparallel fibers (microtubules (Bieling et al. 2010)). In addition to these novel properties, those properties that are shared by prokaryotic filaments also evolved additional layers of regulation. A host of accessory cytoskeletal factors appeared early in eukaryote evolution. For example, microtubule dynamics is regulated by nucleating (gamma-tubulin ring complex [gamma-TuRC]), stabilizing (MAPs), destabilizing (stathmin, katanin), minus-end stabilizing (patronin/ssh4 (Goodwin and Vale 2010)), and plus-end-tracking proteins (+TIPs) (Jiang and Akhmanova 2011). Similarly, actin dynamics is also regulated by a range of accessory factors, mostly representing eukaryotic novelties (Rivero and Cvrcková 2007; Eckert et al. 2011).

The most dramatic innovation in eukaryotes is the use of molecular motors (Vale and Milligan 2000). Kinesins and myosins share a catalytic core that undergoes a conformational change upon nucleotide binding and hydrolysis. This is transmitted by a ‘relay helix’ to the polymer binding sites and the mechanical elements (Kull et al. 1996). Motor proteins are stepping along the filaments using such mechano-chemical cycles. Motors are either nonprocessive or processive, depending whether they perform one or multiple cycles, before detaching from the filament. Processive motion enables long-range transport using one motor protein. A hypothetical evolutionary scheme for the evolution of processive motors from a GTPase is outlined in Figure 5.

**Figure 5.**
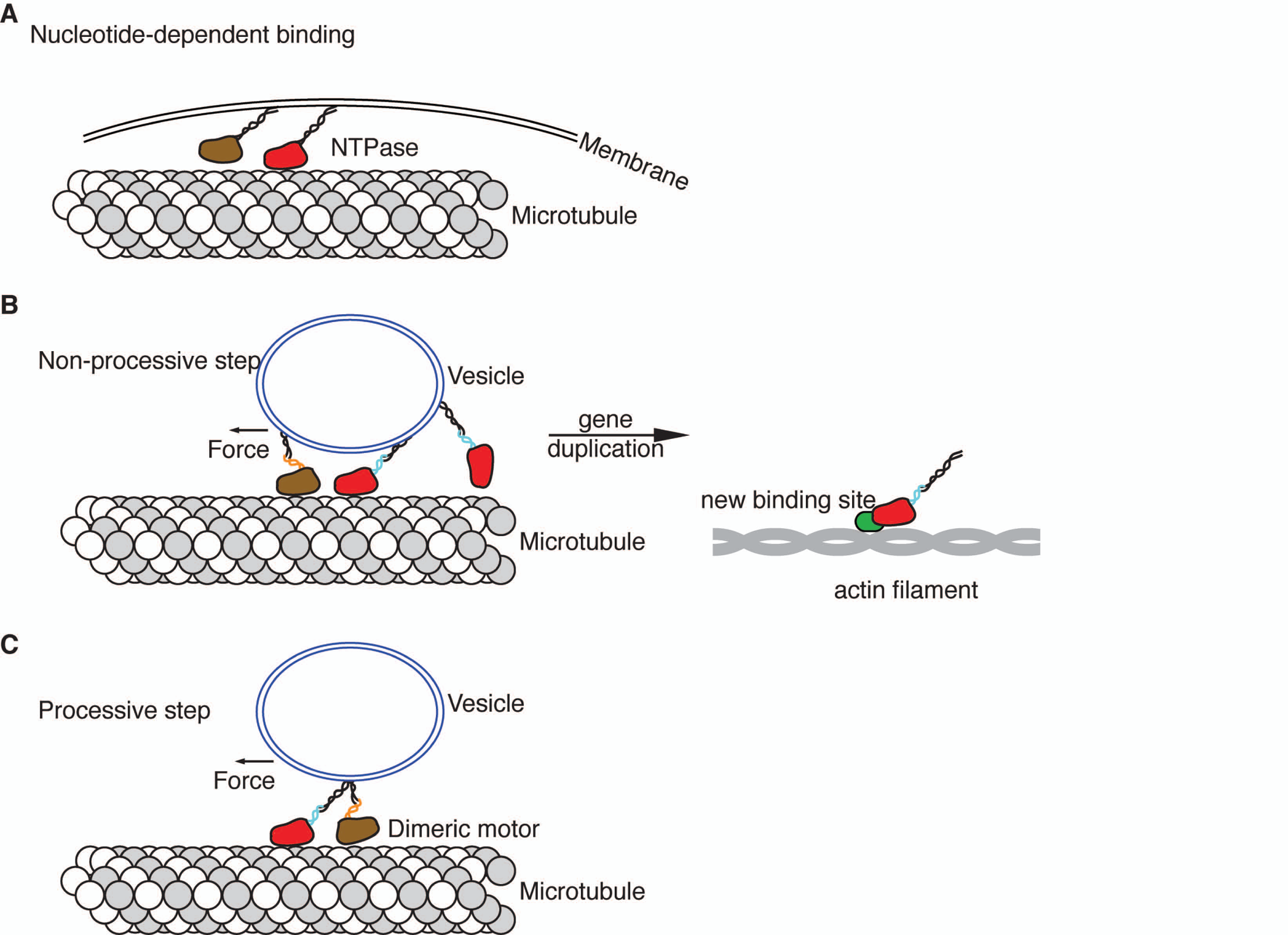
Evolutionary scenario for the origin of processive kinesin and myosin motors. Kinesin and myosin have a common origin, and evolved from a GTPase switch. In the first stage the NTPase bound to the filament in a nucleotide-dependent manner via a short motif connected to the NTPase domain. The NTPase was engaged in other interactions (e.g. membrane binding), and recruited the filament to an organelle. In the next step the mechanical elements evolve that can perform one mechanical cycle following the nucleotide cycle. Motion and dissociation are both coupled to the nucleotide cycle, and are transduced via a relay helix that is conserved between kinesin and myosin. This non-processive motor can now exert force on the bound organelle (e.g. a vesicle). The clustering of several of these motors can move organelles. Monomeric motors may have been non-processive (Berliner et al. 1995), or may have used biased one-dimensional diffusion for processivity (Okada and Hirokawa 1999). Myosin and kinesin probably diverged at such a stage, by the acquisition of a novel filament-binding site and engagement with the second filament type (the direction is unclear). It is unlikely that the common ancestor of kinesin and myosin had a binding surface for both actin and tubulin filaments. Motor dimerization may have evolved to increase the probability of repeated engagement with the filaments. For processivity, the dimensions of the linker had to match the spacing of the accessible binding sites on the filament (80 Å for microtubules, 360 Å for actin filaments). This allowed the filament-dependent coupling of the nucleotide cycles on the two motor heads (the “mechanically controlled access” model) (Vale and Milligan 2000).

The advent of motors added an extra layer of complexity to cytoskeletal dynamics. Motors perform diverse and specialized functions, and are essential for the movement of organelles and complexes (e.g. intraflagellar transport), the establishment of a bipolar spindle, the definition of the cell division plane, cell migration, and cell polarity. For example, specific kinesin families are involved in the regulation of ciliary transport and motility (kinesin 2, 9, 13), and only occur in species with cilia (Wickstead et al. 2010). Others are involved in ciliary length control (Kif19, (Niwa et al. 2012)). Some kinesins regulate different aspects of spindle organization including spindle midzone formation (kinesin-4, Kif14 (Kurasawa et al. 2004; Gruneberg et al. 2006)), alignment on the metaphase plate (Xkid (Antonio et al. 2000)), or centrosome separation during bipolar spindle assembly (Eg5, (Kapitein et al. 2005), Kif15 (Tanenbaum et al. 2009)).

The eukaryotic cytoskeleton forms complex three-dimensional patterns by the dynamic interactions of filaments, motors and accessory proteins in a self-organizing process (Vignaud et al. 2012). The evolution of these new dynamic properties must have been tightly linked to the origin of novel cellular features during eukaryote origins. In the following section I will discuss some of the possible links in the framework of a cell evolutionary scenario.

## Coevolution of a dynamic and scaffolding cytoskeleton with eukaryotic organelles

The origin of the eukaryotic cytoskeleton can be placed into a transition scenario of eukaryote origins. Such scenarios may seem like “just so stories”, but are nevertheless important conceptual frameworks and can identify problems for future research. The first major event that could have precipitated a functional shift in the prokaryotic cytomotive filament systems could have been the loss of a rigid cell wall (Cavalier-Smith 2002). This step is necessary, irrespective of the prokaryotic lineage from which eukaryotes evolved (archaebacteria or the common ancestor of the sister groups archaebacteria and eukaryotes). The loss of the cell wall may have been a dramatic, but not lethal event. The recent discovery of cell division in wall-free bacteria via membrane blebbing and tubulation provides a model for cell division following cell wall loss (Mercier et al. 2013). Importantly, wall-free division is independent of FtsZ, suggesting that in early eukaryote evolution the release of functional constraints may have allowed the rapid functional evolution of FtsZ. Similarly, a rapid shift in prokaryotic actin functions may also have been facilitated by cell-wall loss.

MreB directly contributes to the mechanical integrity of the bacterial cell, independent of its function in directing cell wall synthesis (Wang et al. 2010). This suggests that the mechanical function of the filamentous cytoskeleton may have a prokaryotic origin. Loss of the cell wall may have triggered the elaboration of such a function and led to the evolution of actin networks involved in motility and cytokineses.

An important step in tubulin evolution was the origin of the microtubule, formed by the lateral association of protofilaments. Hollow tubes have higher mechanical rigidity, and could have more efficiently served scaffolding and transport functions. Microtubules may have evolved into the 13-protofilament-form following the origin of the gamma-TuRC complex that helped to fix the number of protofilaments.

A common theme in the evolution of actin and tubulin filaments is the early origin of paralogs involved in nucleating the filaments (Arp2/3 and gamma-tubulin). FtsZ dimers can nucleate FtsZ filaments, and it is conceivable that a gene duplication event allowed the functional separation and streamlining of the filament-forming and nucleating functions for both filament types. The origin of separate nucleating factors allowed a more flexible positioning of nucleating centers, given that these could now be regulated independent of the filaments.

By providing support for membrane transport, the filaments facilitated the evolution of the endomembrane system (Jékely 2003). All endomembranes depend on cytoskeletal factors for their formation and transport. Early membrane dynamics may have evolved to allow endocytic uptake of fluid and particles and to deliver extra membrane to sites of phagocytic uptake. Phagocytosis can proceed even without a dynamic actin cytoskeleton, driven by thermal membrane fluctuations and ligand-receptor bonds that zipper the membrane around a particle. The origin of an actin network could have made this process more efficient by preventing the membrane from moving backwards like a ratchet (Tollis et al. 2010). The origin of phagocytosis could have led to the origin of mitochondria (Cavalier-Smith 2002; Jékely 2007). Energetic arguments seem to favor an early origin of mitochondria (Lane and Martin 2010), however, complex, nucleotide-driven dynamic filament systems are abundant in prokaryotes, and can drive membrane remodeling. For example, overexpresison of the actin-like protein FtsA can lead to the formation of large, protein-coated intracellular membrane tubules (Szwedziak et al. 2012). In addition, amitochondrial eukaryotes can maintain a complex cytoskeleton (e.g. *Trichomonas vaginalis* has 19 kinesin and 41 dynein heavy chains), making the energetic argument for the primacy of mitochondria over phagotrophy less compelling.

The self-organizing properties of the cytoskeleton presumably evolved very early. We know from minimal systems and simulations, that a few components are sufficient to organize complex, dynamic structures, such as spindles, spirals, and aster (Leibler et al. 1997; Surrey 2001; Nédélec 2002; Nédélec et al. 2003). For example, one function of the dynamically unstable microtubule cytoskeleton is to position the nucleus in the cell center by exerting pushing forces on the nucleus (Tran et al. 2001). This process contributes to the spatial organization of the cell (e.g. by determining the cell division plane). Centre-positioning can also work *in vitro* with a minimal system of dynamic microtubules, even in the absence of motor proteins (Holy et al. 1997).

These examples illustrate that we have a growing understanding of the self-organization of dynamic cytoskeletal structures of various shapes and functions. In future studies, this knowledge could be combined with comparative genomic reconstructions to study ‘alternative cytoskeletal landscapes’ in different eukaryotic lineages (Dawson and Paredez 2013), and to reconstruct the stepwise assembly of these self organizing structures during the origin of eukaryotes.

## Concluding remarks

The complex self-organizing properties of the cytoskeleton set it apart from other cellular systems, such as large macromolecular assemblies or metabolic pathways. This means that it is difficult to deduce what effects the addition or loss of one component might have had on the systems-level properties. This is in contrast to metabolic pathways, where evolutionary changes can be efficiently modeled using flux-balance analysis of the entire metabolic network of a cell (Pal et al. 2005). A similar analysis is not yet feasible for the entire cytoskeletal network. However, it would now be possible to study the evolution of sub-systems from a systems perspective. Consider the mitotic spindle. We have a good understanding of how the antiparallel microtubule arrays overlapping at their plus ends form in a dynamic process involving an interplay of microtubule growth and shrinkage, motor activity, and proteins binding specifically to the overlap region (Janson et al. 2007). The emergence of a dynamic bipolar spindle can also be captured in computer simulations (Nédélec 2002). In evolutionary models, one would have to consider a succession of states following the gradual change of activities or addition of components. There are at least five ancestral kinesin families involved in mitosis (Wickstead et al. 2010). How did mitosis work when there was only one kinesin, in a stem eukaryote?

The origin of axonemal motility, involving microtubule doublets and at least seven ancestral axonemal dynein families (Wickstead and Gull 2007), represents a similar problem. What was the beat pattern of the proto-cilium like, with only one axonemal dynein? How did it change when inner-arm and outer-arm dyneins diverged? The origin of lamellipodial motility and phagocytosis could also be best addressed by focusing on minimal systems that allow the formation of membrane protrusions supported by an actin network (Gordon et al. 2012; Vignaud et al. 2012). Only a combination of mutant studies (Mitchell and Kang 1991), in vitro reconstituted systems (Takada and Kamiya 1994), comparative genomics (Wickstead and Gull 2007), and computer simulations (Brokaw 2004; Tollis et al. 2010) could answer these questions.

## Acknowledgments

I thank David R. Mitchell and Elizabeth Williams for their comments on the manuscript. The research leading to these results received funding from the European Research Council under the European Union's Seventh Framework Programme (FP7/2007-2013)/European Research Council Grant Agreement 260821.

